# Single strain control of microbial consortia

**DOI:** 10.1101/2019.12.23.887331

**Authors:** Alex J.H. Fedorec, Behzad D. Karkaria, Michael Sulu, Chris P Barnes

## Abstract

The scale of the biological systems we can engineer is limited by the burden that host cells can bear. Division-of-labour can spread that burden across a community of cells but competitive exclusion inevitably leads to the removal of less fit community members over time. Here, we leverage amensalism and competitive exclusion to stabilise multi-species communities by engineering a strain of *Escherichia coli* which secretes a toxin in response to competition. We show mathematically and experimentally that such a system can produce stable populations with a composition that is tunable by easily controllable parameters. This is the first system to use competitive exclusion to create a stable two-species consortia and the first to only require the engineering of a single strain.

## Introduction

Techniques for the assembly [8] and synthesis [18] of DNA sequences has enabled the construction of chromosome scale [1,43], synthetic biological systems. Recent efforts to systematically characterise genetic “parts” [12, 46] and develop software tools to produce DNA sequences from function specification[30] are enabling synthetic biological systems in which large numbers of regulatory proteins and promoters are involved. The production of each of these proteins sequesters resources that would otherwise be used by the host organism for growth and therefore leads to a reduction in growth rate [22, 9]. This provides a selection advantage to loss-of-function mutants that arise, suggesting that the functional period for the increasingly complex synthetic biological systems of the future will become ever more ephemeral.

Attempts have been made to minimise burden [9, 10], make mutation deadly [5], or periodically displace mutating populations with a functioning population [26]. Over the past decade there have been attempts at division-of-labour in which a system is split into subcomponents and distributed into specialised subpopulations of cells [32, 40]. This minimises the burden that is placed on individual cells which reduces, but does not remove, the selective advantage of loss-of-function mutations In addition, the creation of synthetic communities allows the diversification and compartmentalisation of functions, modularisation, spatio-temporal control and mechanisms for biosafety [20].

The fundamental challenge with constructing such heterogeneous communities is the principle of competitive exclusion, which states that two species competing within the same niche cannot coexist [16]. The principle should, perhaps, include the caveat: “in stable environments and in the absence of other interactions”, which may help to explain supposed deviations such as the “paradox of the plankton” [19]. Wild bacteria live in complex communities [38] with mutualistic and competitive interactions producing complex dynamics[17]. Previous attempts to design synthetic microbial communities have relied on spatial segregation [21] or mutualism [37] to maintain multiple sub-populations The control of the density of monoculture has been achieved through the use of quorum sensing to control self-killing [47] and recently this has been extended to a two species system [35]. More complex predator-prey systems have also been developed that produce oscillatory populations of two strains [3]. These systems involve the engineering of all strains within the community. However, this requirement may not be desirable in industrial settings and is clearly not possible when working in natural environments such as the human gastrointestinal tract.

In order to control a community through a single constituent, we require a mechanism that allows control of the growth rate of one or more competitors at a distance. Here we suggest bacteriocins, secreted anti-microbial peptides, as such a mechanism and detail the construction and characterisation of a control system that uses them. A wide range of bacteriocins are produced in natural microbial communities such as the human intestinal microbiota [13] where they play an important role in niche competition [23]. We have previously demonstrated the ability of the bacteriocin microcin-V to improve plasmid maintenance [14]; a challenge which includes preventing competitive exclusion. Further, gram positive bacteriocins have been used to produce commensal and amensal interactions along with all pairwise combinations of the two [24]. By limiting ourselves to the engineering of a single strain, this system could be repurposed for many applications by only selecting the bacteriocins appropriate for the target niche.

## Results

### Bacteriocins enable population control at a distance

We engineered an *E. coli* JW3910 strain [2] to express microcin-V and its immunity gene from a plasmid (Methods). In addition, the plasmid carries the transport mechanism to secrete the fully formed bacteriocin to kill a competitor strain, *E. coli* MG1655, Figure 1A. The engineered strain also constitutively expresses mCherry in order to produce a burden, which reduces the growth rate and allows the separation of the engineered and competitor strains in the microplate reader [33]. In order to demonstrate that the burden placed on the engineered strain leads to competitive exclusion, we mutated the start codon of the *cvaC* bacteriocin gene [15], rendering it incapable of killing the competitor. As the two subpopulations grow, the faster growing competitor quickly dominates while growth substrate is abundant in the first two hours, Figure 1B. As the growth substrate begins to run out, the rate of competitive exclusion reduces but still occurs while the population is growing. The strain with the intact bacteriocin is capable of killing the competitor and, over time, dominates the community, Figure 1B.

**Figure 1:**
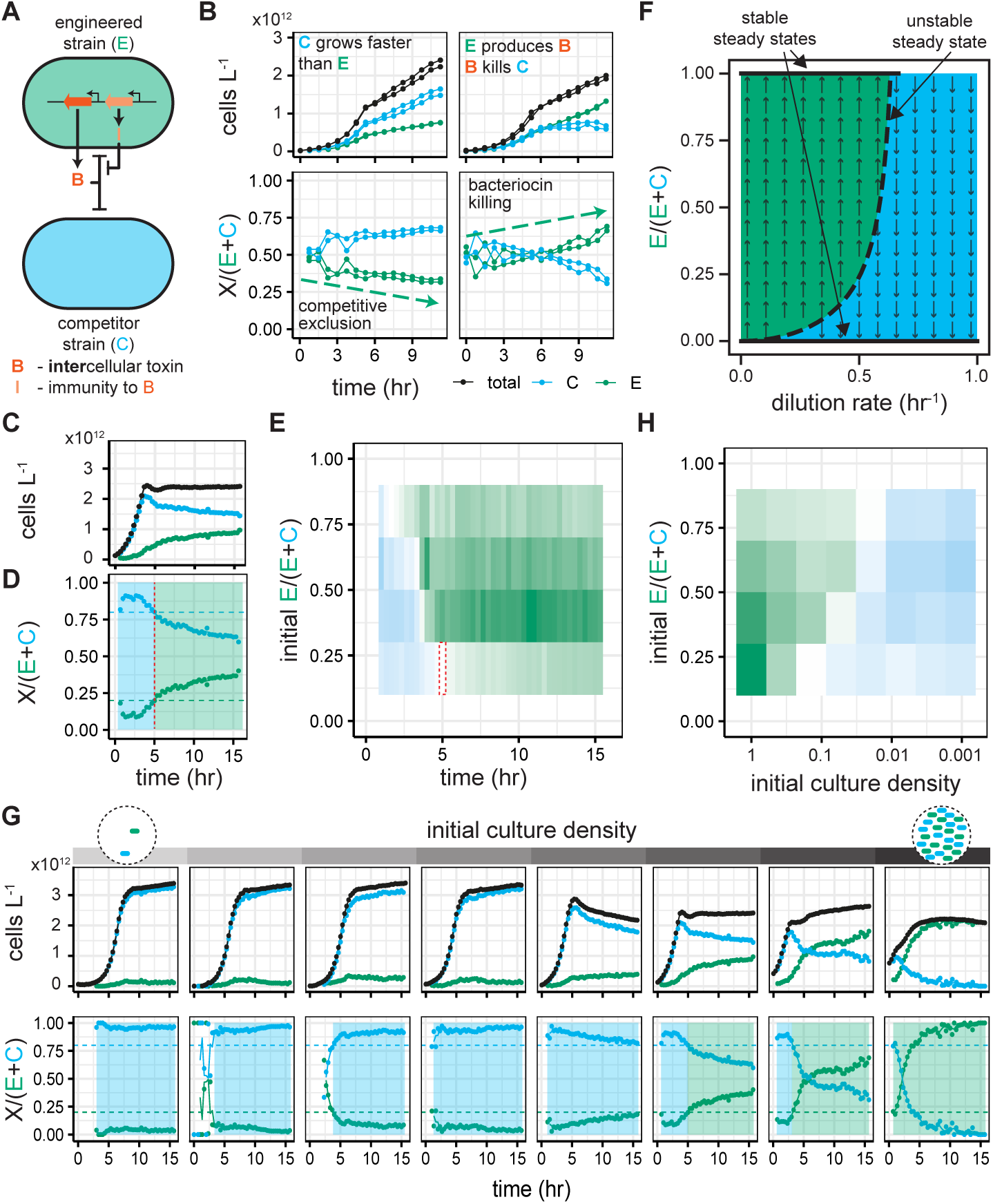
Bacteriocins can be used to overcome competitive exclusion. (A) The engineered strain produces a bacteriocin and corresponding immunity protein. Once secreted, the bacteriocin kills the competing strain. (B) With an inactivated bacteriocin, the faster growing competitor excludes the engineered strain. However, secretion of an active bacteriocin leads to killing of the competitor and domination by the engineered strain. (C) The two strains are co-cultured in a microtitre plate and the subpopulation fractions are determined using fluorescent proteins in the engineered strain and the flopR software [33], allowing us to track the population dynamics over time. (D) The blue shaded area shows the period during which the competitor ratio is above its initial ratio; the green shaded area shows the same for the engineered strain. The red dashed line indicates the timepoint at which the change from competitor to engineered strain dominance occurs. (E) Graph showing the “winning” strain at each time point of co-cultures with different initial-ratios. The colour and saturation of each block is given by the difference between the initial strain ratio and the ratio at that timepoint. The dashed red box shows the point in (D) at which the engineered strain begins to win. (F) Analysis of the mathematical model with varying dilution rate. Solid black lines show the stable steady states (extinction of one or other strain), the dashed black line indicates an unstable steady state. The arrows show the direction of population change either side of the unstable manifold. (G) Dilution rate is approximated by varying the initial density of the co-cultures in the microtitre plate; lower initial density approximating faster dilution rates. We see the same switch, from competitive exclusion to bacteriocin killing, predicted by the model. (H) Graph of the “winning”strain after 5 hours of growth in co-culture, over a range of initial population ratios.

When we look more closely at the population dynamics of a co-culture, with bacteriocin production over time, we observe an initial phase of competitive exclusion before the killing of the competitor occurs, Figure 1C & D. The time it takes before the engineered strain outcompetes the competitor (the point at which the proportion of the population constituted by the engineered strain is greater than its initial proportion) is related to the initial population ratio, Figure 1E. A lower initial ratio of engineered strain leads to a longer period in which the competitor is winning, and *vice versa*. This suggests that a simple strategy to control these competing populations in a batch environment would be to passage more frequently if the engineered strain is dominating or less frequently when the competitor dominates. One can view this as the competitor strain being fitter in environments with a high dilution frequency and the engineered strain being fitter in low dilution frequency environments.

## Restricting growth favours bacteriocin producers over fast growers

Using a simple mathematical model (Supplementary Information) we are able to simulate growth dynamics and bacteriocin production in a chemostat environment. Analysis of our model shows that at lower dilution rates there are two stable steady states, in which one or the other of the two strains goes extinct, and an unstable coexistence steady state, Figure 1F. At higher dilution rates, only the competitor strain is able to dominate and with no dilution only the engineered strain will survive. The green region shows the area in which the community will tend towards engineered strain dominance over time; the blue region shows the area in which the competitor will dominate over time. This reinforces our previous statement that, with control of dilution rate, one could switch between states in which the engineered strain or the competitor strain dominates based on the current community composition. A similar approach has previously been demonstrated using nutrient control rather than dilution rate [42].

The dilution rate in a chemostat directly affects the density of the bacteria and their growth rates via substrate concentration. In addition, the dilution rate will also affect the concentration of bacteriocin that is able to accumulate. We explore how these impact community dynamics by serially diluting a co-culture with fresh media to produce a range of initial culture densities, Figure 1G. In the most dilute cultures, the initial concentration of bacteriocin is very low and the growth substrate is plentiful allowing the competitor strain to dominate the population before the engineered strain is able to get a foothold. As the initial density increases so does the initial bacteriocin concentration and the number of engineered cells secreting bacteriocin. At the same, the initial substrate concentration is reduced, narrowing the window over which the competitor can outcompete the engineered strain. This combination leads to a shortening of the period of competitive exclusion and an increase in the rate at which the engineered strain takes over, Figure 1G. The same experiment is performed with varying initial co-culture ratios, Figure 1H and Supplementary Figure 1. We see that, as the initial density increases, the system moves from favouring the competitor to favouring the engineered strain, just as we showed by changing the dilution rate with our mathematical model in Figure 1F.

The death rate of the competitor strain reduces after the cultures reach a peak density; this is most notable in the penultimate initial density shown in Figure 1G. This demonstrates a potential point of deviation between the mathematical model, in which cells are in a chemostat and therefore growing exponentially, and our microplate reader experiments in which stationary phase is reached. The 5 hour sample timepoint chosen for Figure 1H and later figures is approximately at the end of the exponential growth phase and before the change in killing rate occurs.

### Flipping competitive advantage via an exogenous inducer

For some applications, total control of the environmental parameters such as dilution rate is unrealistic. As such, we need another mechanism for switching state from one strain to the other dominating. We can achieve this through control of the production rate of the bacteriocin, Figure 2A. To implement such a system, we constructed a biological circuit, split across two plasmids, in which expression of microcin-V is repressed by TetR, the expression of which is induced by *N*-3-oxohexanoyl-homoserine lactone (3OC6-HSL), Figure 2B. Using a agar spot inhibition assay, we demonstrate that we can exogenously control the killing of a competitor in a dose dependent manner, Figure 2C & D; which to our knowledge is the first demonstration of dose dependent killing with bacteriocins. Our mathematical model demonstrates that, at lower rates of bacteriocin production, the killing of the competitor is reduced, allowing it to dominate, whereas at higher bacteriocin production rates the engineered strain dominates, Figure 2E.

**Figure 2:**
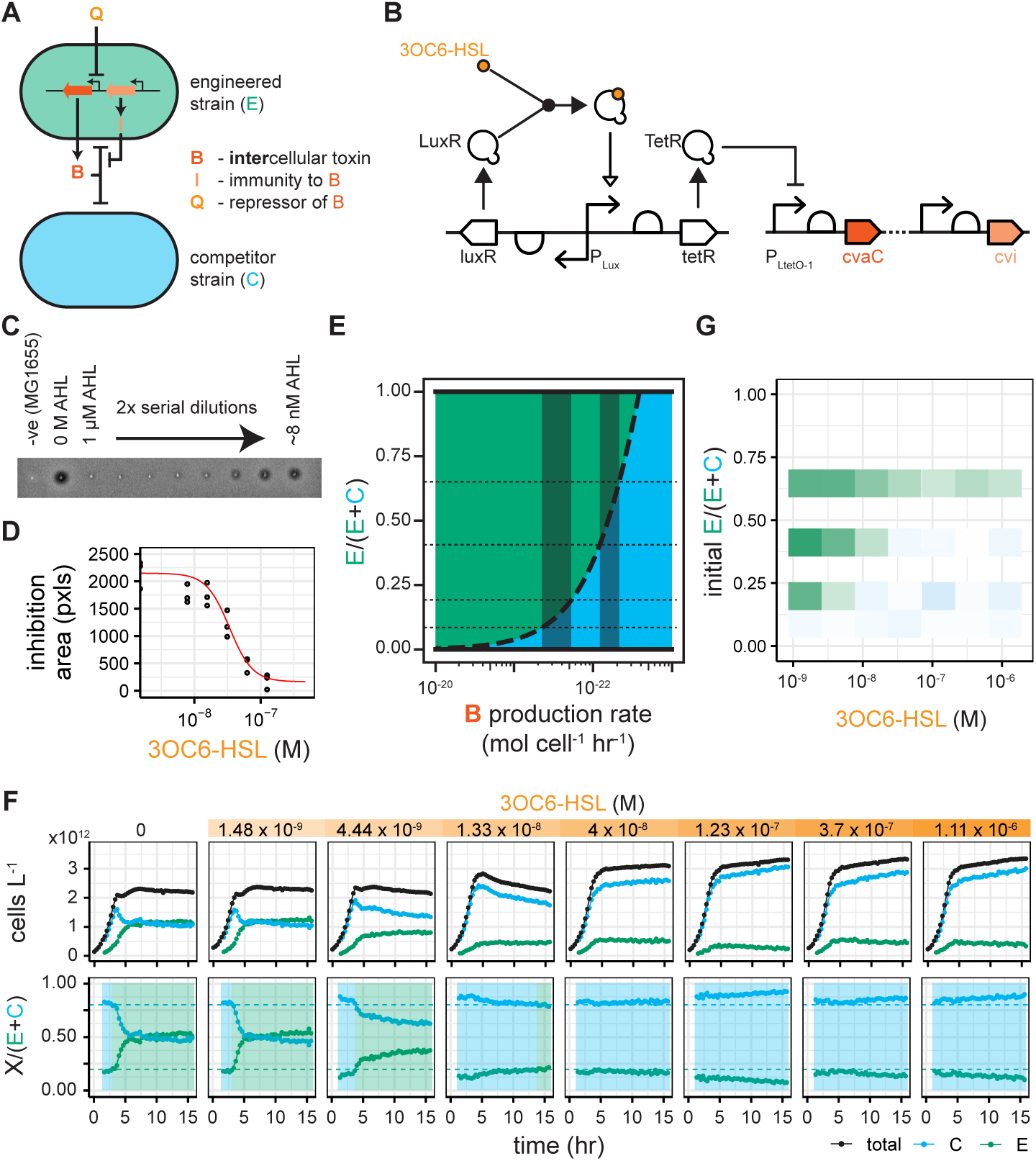
Exogenous control of bacteriocin production. (A) The expression of bacteriocin can be controlled by the addition of a quorum molecule (Q = 3OC6-HSL) into the environment. (B) The system is spread across two plasmids. 3OC6-HSL binds to LuxR, *luxR*-AHL induces the expression of TetR which represses bacteriocin expression. (C) Conditioned media from the growth of the engineered strain in a range of 3OC6-HSL concentrations was spotted onto a lawn of EcM-Gm-CFP (Supplementary Information) leading to zones of inhibition from bacteriocin killing. (D) The inhibition areas are quantified using an image processing pipeline, and used to fit a Hill function for bacteriocin expression. (E) Analysis of the mathematical model with varying bacteriocin expression rate. n.b. the *x*-axis is reversed to mimic the direction of 3OC6-HSL control i.e. high bacteriocin production rate on the left, low bacteriocin production rate on the right. The dotted horizontal lines mark the initial population ratios from (G). The shaded regions show the possible regions of maximum and minimum bacteriocin production rate based on (G). (F) Dynamics of co-cultures grown in different concentrations of 3OC6-HSL. (G) A graph of the “winning” strain after 5 hours of co-culture, with varying 3OC6-HSL concentration and across several initial population ratios. Mapping these results onto (E) shows the possible regions of minimal and maximal bacteriocin expression rate within which our system is operating.

Communities of the engineered and competitor strains were grown in various concentrations of 3OC6-HSL and tracked in the microplate reader, Figure 2F and Supplementary Figure 2. We see that, as the concentration of 3OC6-HSL increases, the killing of the competitor by the engineered strain is relieved and the system moves from engineered strain domination to competitor domination. This is reflected in the changes that we see after 5 hours of competition, Figure 2G. However, if the population starts with a low proportion of engineered cells, even at the lowest concentration of 3OC6-HSL, the engineered cells are unable to compete. Similarly, if the engineered cell population constitutes a high proportion of the initial culture the competitor is unable to compete, regardless of the amount of 3OC6-HSL. This means that we can only control which subpopulation dominates within a narrow range of consortia ratios; outside of this range we cannot prevent the eventual extinction of one of the strains. The former scenario could be rectified by increasing the initial culture density or running for a longer period, as we showed in Figure 1. However, the latter case is rectified with the inverse solution; lower initial density of shorter passage length. This arises from the fact that, in the system we have built, the maximal fold change in expression level of the bacteriocin is ∼ 10, as shown in Figure 2D. This is further reflected in Figure 2E where, using the population ratios at which we have data, we have overlaid the possible regions for minimal and maximal bacteriocin production rate; the extreme ends of the regions is ∼ 10 fold difference in bacteriocin expression. Ultimately, increasing the fold change in bacteriocin expression will increase the range of consortia composition over which our system can operate.

### Fully autonomous community regulation

The use of the quorum sensing molecule 3OC6-HSL to control the bacteriocin allows us to build a system in which the density of the engineered strain dictates the expression level. Using cell density to control system response affords the creation of an autonomous population control system in which actions are taken by cells dependent on the state of the community rather than through external control. This approach has previously been taken to control homogeneous [4] and heterogeneous populations [35] with intracellular toxins. Here we add the ability to tune the rate of production of the 3OC6-HSL through arabinose induction of the quorum molecule synthase, LuxI, Figure 3A. This system should enable the engineered strain to sense competitive exclusion through a decrease in the concentration of 3OC6-HSL. In response, the engineered strain produces and secretes the bacteriocin, increasing its fitness. As the population of the engineered strain increases, so to does the concentration of 3OC6-HSL, leading to repression of the bacteriocin and allowing the competitor to grow again. In this way the engineered strain can dynamically balance the population ratios, Figure 3B. The system was constructed using a modular approach that allowed us to test the function of each component as we progressed, Figure 3C, and enabled us to use modelling during the construction and characterisation to determine whether the system, as constructed, could achieve the stable co-existence that we desired. Fluorescent proteins were cloned downstream of each promoter for characterisation and flow cytometry was used to quantify expression levels. We used a Bayesian approach to fit Hill functions to the flow cytometry data, Figure 3D. The copy number of the plasmid bearing the control elements (arabinose inducible LuxI and 3OC6-HSL inducible TetR) was explored (Supplementary Figure 4) to ensure the maximal expression range of TetR, inferred from GFP expression, under varying arabinose concentrations without overburdening the host cells.

**Figure 3:**
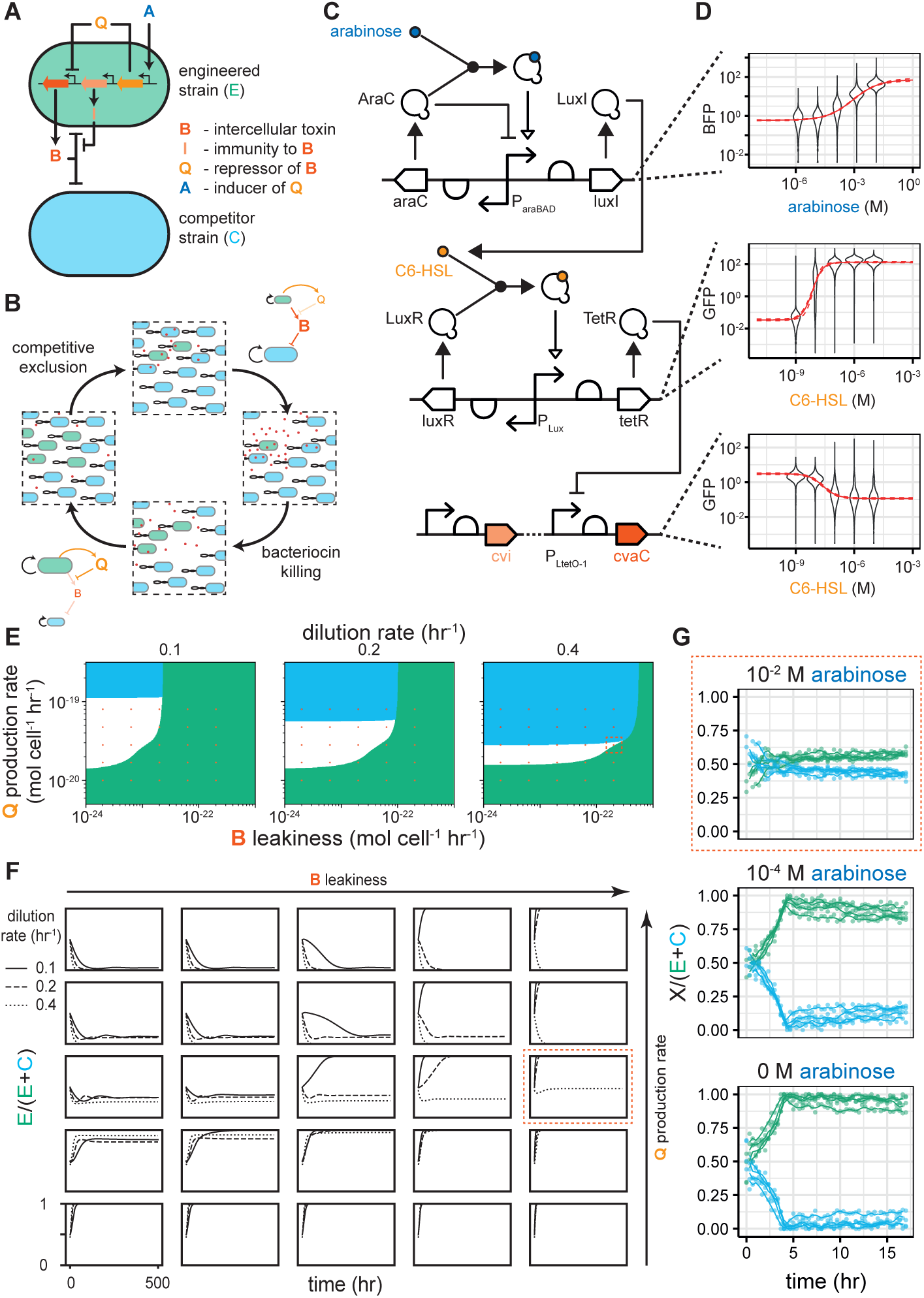
Autonomous control of the community through quorum sensing. (A) The engineered strain produces its own quorum molecule. The rate of production of the quorum molecule synthase is tunable by arabinose induction. (B) Bacteriocin production is turned off when the engineered population density is high, allowing the competitor to grow, and competitive exclusion to complete the cycle. (C) The system was constructed in three parts: arabinose inducible expression of LuxI, 3OC6-HSL inducible expression of TetR, and TetR repressible expression of microcin-V. (D) Each part was characterised using fluorescent reporter proteins (GFP or BFP) and flow cytometry. Hill functions were fitted (violins show all events from 3 replicates) using a Bayesian method (solid line = mean prediction, dashed lines = 95% credible region in the mean). (E) Steady state analysis and (F) simulation with varying quorum molecule production rate and bacteriocin expression leakiness, at three different dilution rates, shows parameter values at which the engineered (green) or competitor strain (blue) dominates. The white regions show areas of stable coexistence. The red circles indicate the position in parameter space at which the timecourses in (F) are simulated. The dashed red boxes (in E, F and G) indicate the region in which coexistence in the microplate reader has been acheived. (G) Population dynamics of co-cultures at different arabinose concentrations. At 10 mM arabinose, coexistence is observed across the timecourse at the population ratio predicted by the model. Lower arabinose concentrations lead to dominance of the engineered strain.

**Figure 4:**
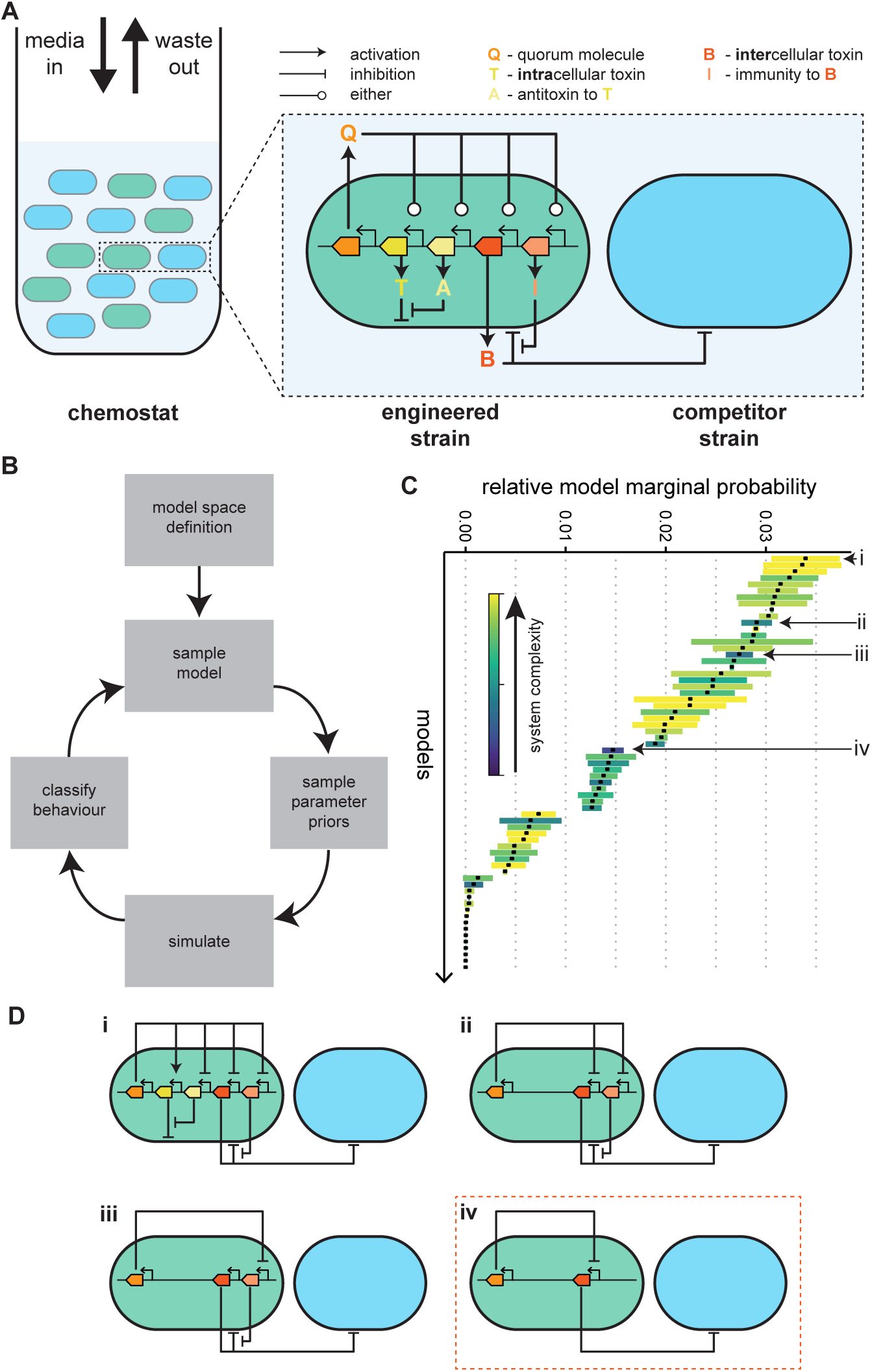
Exploring the space of possible models of population control to find the best system. (A) Schematic of all possible components for single strain engineering. The engineered strain can produce an intercellular toxin and its antitoxin, an intracellular toxin and its antitoxin, and a substrate to enhance growth. All of these can be positively or negatively regulated by a self-produced quorum molecule, allowing density dependent control of the engineered cell’s actions. (B) Once the model space has been defined, we explore it using ABC SMC to determine which models with which parameters are capable of producing stable coexisting populations. (C) The ordered model marginal posterior probabilities. The long tail of extinct models has been trimmed. The boxes for each model are coloured according to the model’s complexity; models with a larger number of parameters are closer to the yellow and fewer parameters are close to blue. (D) The best system (i) requires control over the expression of four genes. Systems that don’t require intracellular toxin or immunity perform well for their level of complexity (ii - iv). In fact the simplest system able to produce stable coexistence is the system that we have developed (iv).

A steady state analysis of this system shows regions of stable coexistence which vary in size depending on the dilution rate of the environment, Figure 3E. The extent to which we can repress the bacteriocin also has an effect, with leakier systems narrowing the range of 3OC6-HSL production within which steady states are achievable. If the system is too leaky, the engineered strain always dominates. Importantly, changing the production rate of 3OC6-HSL, which we can do through arabinose induction of LuxI, allows us to move the system into a region with stable coexistence. In addition, we can tune the population ratio within the stable coexistence region by using the 3OC6-HSL production rate, with a higher rate resulting in a lower fraction of engineered strain in the community, Figure 3F. Our estimates of bacteriocin leakiness from Figure 2D, suggest that stable coexistence is theoretically possible with a high dilution rate. However, the region for 3OC6-HSL production rate is very small, suggesting achieving this would be difficult. We ran co-cultures in a microplate reader with increasing amounts of arabinose, at a starting density which, based on results in Figure 1, we believe to produce dynamics akin to a dilution rate of 0.4 hr^−1^, Figure 3G. Without arabinose and at low concentration, the engineered strain dominates the culture, as predicted. However, at 10 mM arabinose, we see coexistence for the length of the experiment, mimicking the dynamics seen in the simulation. It should be noted that the plate reader is a limited approximation of the chemostat environment assumed in our mathematical model. Despite this, we have managed to capture the predicted behaviour.

### Identifying robust models for consortia control

Alongside our intercellular toxin mechanism for controlling the fitness of a population at a distance, we can also incorporate intracellular toxins which have been used before for population control. In addition to the expression of these toxins, it has also been suggested that their respective immunity genes can be regulated to change the viability of the engineered strain [29]. One could additionally include the production of a growth enhancing substrate, but we neglect this as it would either require engineering, or careful selection, of the competitor strain. The expression of all of these molecules can be constitutive or under control of a quorum molecule also expressed by the engineered strain, Figure 4A.

By defining a set of expressible parts that can be assigned to a strain, we can produce a model space with the stipulation that only a single strain is engineered. In this case the space consists of 132 unique systems for which we can compare their ability to produce stable co-existing communities. From a relatively small number of available parts, we have a large number of possible models to choose from. In order to narrow our search to a smaller set of candidate models we perform model selection using approximate Bayesian computation sequential Monte Carlo (ABC SMC), Figure 4B. This approach allows us to approximate model and parameter posterior probabilities by random sampling and weight assignment through a series of intermediate distributions [41, 45]. The output of ABC SMC is an approximation of the posterior distribution of models and the posterior distribution of parameters for each model. The final model posterior distribution indicates which models have the highest probability of producing the objective behaviour – in this case coexisting communities at steady state – while also accounting for system complexity (Occam’s razor), Figure 4C. All of the top performing systems use bacteriocin. Indeed, the best system without bacteriocins is ranked 75th. The best system overall requires the control of all four of the possible genes; the bacteriocin, immunity and antitoxin are repressed by the quorum molecule, while the intracellular toxin is induced by it, Figure 4Di.

The systems without the intracellular toxin and antitoxin perform particularly well when one considers their relative simplicity. The best of these uses the quorum molecule to turn off expression of both the bacteriocin and immunity, Figure 4Dii. Intuitively, this leads to less killing of the competitor at higher densities, but also increased susceptibility of the engineered strain. A slightly simpler system uses the quorum molecule only to repress the immunity gene, Figure 4Diii. The simplest system to robustly achieve stable coexisting populations is the system that we have built, requiring just the bacteriocin to be repressed by the quorum molecule, Figure 4Div. The immunity gene is expressed in this system but, as its constitutive expression level doesn’t need tuning, we consider this system simpler than Figure 4Diii.

None of the previously described systems [35, 29, 3] are able to produce stable communities. However, if we set our objective to co-existence, with both populations above a defined threshold, rather than stable communities, the other systems can achieve the goal, Supplementary Figure 5. The theoretical system proposed by McCardell *et al*. performs as well as the system we have constructed. Just as for stable communities, the addition of quorum control over immunity expression, as in Figure 4Dii, improves robustness further.

These results demonstrate that we have constructed a ‘best in class’ system using only a limited number of parts, which is expected to be more robust than existing systems. It also provides a path to implementing more robust systems in the future.

## Discussion

We have demonstrated that bacteriocins can be used to control the relative fitness of strains in competition with one another. The environment is a key determinant of strain fitness. We have shown that dilution rate alone can be used to switch between the domination of a fast growing strain and an amensal strain. Although microplate reader assays, in which cultures grow to saturation on a limited substrate, do not fully reflect the chemostat environment which we have modelled, our experimental results closely follow the model predictions. The inclusion of exogenous control through 3OC6-HSL allowed us to demonstrate, for the first time, the dose dependent expression of a bacteriocin. However, we were only able to achieve a ten-fold change in expression, while the modelling results suggest an increase to 100-fold would enable a system with true switching ability regardless of the community composition. Using 3OC6-HSL also enabled us to extend the system to a fully autonomous one in which population density provided the switch from engineered strain to competitor strain domination. The dynamic control of intracellular toxins through quorum sensing has been demonstrated before for community control [47, 35]. However, self-limiting systems are susceptible to loss-of-function mutations and require the engineered strain to grow faster than the competitor, or control of both strains. Although intercellular toxins have been used for synthetic ecologies [24], autonomous control has never before been demonstrated. Further, the embrace of competitive exclusion has not been explicit in any synthetic community control system. Uniquely, this allows the construction of synthetic communities while only requiring the engineering of a single strain, a feature that enables applications of synthetic biological systems in which native microbiomes will be competing against the engineered population. In the shorter term, this system can be used in industrial biotechnology for the control of communities in bioreactors in which division-of-labour is desirable to prevent burden. Our ABC SMC model exploration has demonstrated that the system we have implemented is the simplest system to produce stable coexistence of competing populations; performing better than the intracellular toxin approaches used to date. Further, it has suggested designs to upgrade the system in order to make the community control more robust. This method of model exploration could easily be extended to explore a model space in which the requirement for single strain engineering is dropped or other control mechanisms, such as substrate cross-feeding, are introduced.

The last decade has seen synthetic biology recognise the importance of context, be it compositional, intracellular or environmental [7, 6]. However, community context has largely been neglected, with the consequence that we have limited ourselves to building systems of single, homogeneous populations that are only capable of functioning in controlled environments. Contending with intracellular context has required the use of feedback to take into account the cell’s response to our demands [36, 10]. By embracing competitive exclusion we have shown that feedback is also a crucial component in the construction of stable synthetic microbial communities and have demonstrated for the first time how to work with community context rather than against it.

## Methods

### Strains and Plasmids

A number of plasmids were created for this work, listed in Table 1. All plasmid sequences were confirmed with Sanger sequencing and the plasmid maps are available online. Where the created plasmids have used SEVA vectors [28], the SEVA naming convention for antibiotic resistance and origin of replication has been adhered to, though the “SEVA” prefix has been removed as the plasmids don’t necessarily adhere to the SEVA standards. The SEVA convention for naming of the “cargo” region has not been adhered to as the extensibility of this naming convention is limited.

**Table 1:**
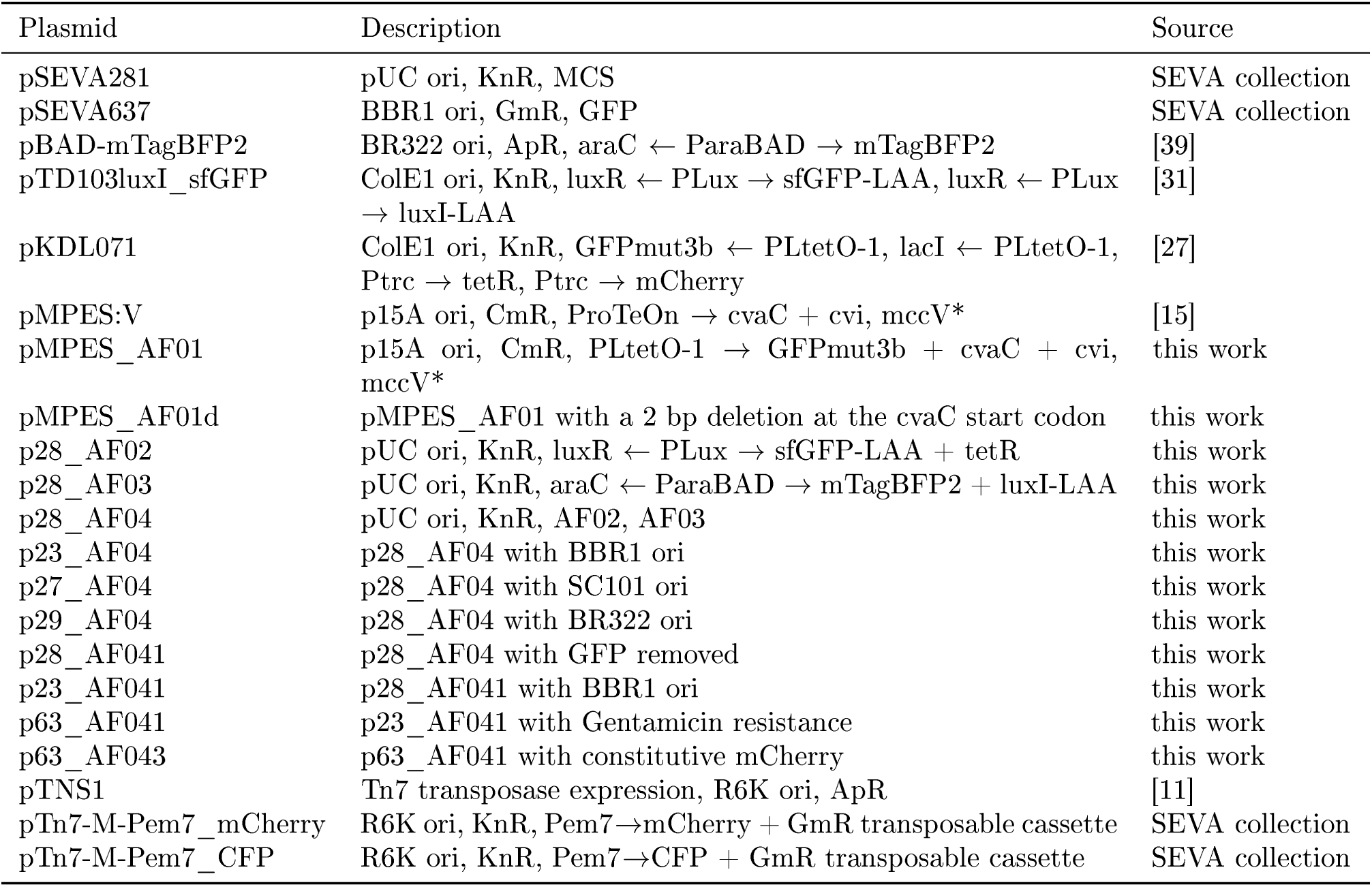
Plasmids used in this work.

pBAD-mTagBFP was a gift from Vladislav Verkhusha (Addgene plasmid 34632; http://n2t.net/addgene:34632 ; RRID: Addgene_34632). pTD103luxI_sfGFP was a gift from Jeff Hasty (Addgene plasmid 48885 ; http://n2t.net/addgene:48885 ; RRID Addgene_ 48885) pTNS1 was a gift from Herbert Schweizer (Addgene plasmid 64967 ; http://n2t.net/addgene:64967 ; RRID: Addgene_ 64967)

The bacterial strains used in this work are detailed in Table 2. Two strains were produced with chromosomal integration of Gentamicin resistance and fluorescence (CFP or mCherry) using the mini-Tn7 transposon system [25] with plasmids from the SEVA collection [28]. Electrocompetent cells were produced using the mannitol-glycerol step protocol [44].

**Table 2:** Bacterial strains used in this work.

| Strain | Description | Source |
| --- | --- | --- |
| *E. coli* NEB 5-alpha |  | New England Bio-Labs |
| *E. coli* JW3910 | Keio collection methionine auxotroph [2] | Horizon Discovery |
| *E. coli* MG1655 |  | Prof. John Ward (UCL) |
| EcM-Gm-CFP | *E. coli* MG1655 with chromosomal Gentamicin resistance and CFP | this work |

### Plate reader competition assays

Overnight cultures of the relevant strains (Competitor = EcM-Gm-CFP, Engineered strain = *E. coli* JW3910: p63_AF043 + pMPES_AF01) were grown, from single colonies picked from selective LB agar plates, in 5 mL of supplemented M9 media(0.4% glycerol 0.2% casamino acids) containing all necessary antibiotics. The overnight cultures were diluted 1:1000 into fresh M9 media containing 10 μg per mL Gentamicin and relevant inducers and grown for 6 hours. The cultures were diluted to an OD (700 nm) of 0.1 using fresh M9 media containing 10 μg per mL Gentamicin and relevant inducers. The cultures were then mixed at the specified ratio. 10 μL of the culture was inoculated into 115 μL of media containing Gentamicin and relevant inducers in the wells of a 96 well microtitre plate and the plate was covered with a Breathe-Easy sealing membrane. The plate was placed in a plate reader (Tecan Spark) and grown for shown number of hours at 37°C with continuous double orbital shaking (2 mm, 150 rpm). Measurements of absorbance (600 nm and 700 nm) and fluorescence (CFP: ex = 430 nm, em = 490 nm, GFP: ex = 485 nm, em = 530 nm, mCherry: ex = 575 nm, em = 620 nm) were taken every 20 minutes. Data was analysed using flopR [33] to produce population and subpopulation curves.

### Deviations from the above

Figure 1B: Wells in the the microtitre plate were inoculated with 200 μL of culture and a magnetic, automatically removable, plastic lid was used to cover the plate. Measurements were taken every 45 minutes. Figure 1C, D, F and G: Rather than inoculating 115 μL of media in each well with 10 μL of culture, the first row of the plate was inoculated with the culture at 01. OD_700_. Subsequent rows were serial dilutions with a dilution factor of 0.375 i.e. by the 8th row the culture was at approximately 1000th the density of the culture in the first row.GFP fluorescence was measured with emission = 535 nm and mCherry fluorescence was measured with excitation = 561 nm. Figure 2E and F: The large amount of time taken to dilute culture to 0.1 OD_700_ meant that by the time cultures were mixed, some cultures, notably the competitor cultures, were no longer at the desired OD. As such, initial ratios were not simply the mixing ratios. We instead determine the initial population ratios from the plate reader data.

### Agar Plate Spot Inhibition Assay

Cultures of the bacteriocin producing strain (*E. coli* MG1655: p23_AF041 + pMPES_AF01) were grown overnight in supplemented M9 media without antibiotics but with the recorded concentration of inducer. A culture of a bacteriocin sensitive strain (EcM-Gm-CFP) was also grown overnight. A one-well plate was filled with 30 mL of 1% LB agar with Gentamicin and allowed to set. 10 mL of 0.5% LB agar with Gentamicin was inoculated with 100 μL of the overnight culture of the bacteriocin sensitive strain, poured over the one-well plate and allowed to set for 1 hour. The overnight cultures for sampling were spun down at 4,000 rpm for 10 mintues and 3 μL of supernatant from each culture was spotted on to the surface of the lawn and the plate was allowed to dry for a further hour. The plate was placed in an incubator at 37°C for 20 hours. The plate was visualised in a BioRad GelDoc using the epi-white light source.

An image processing pipeline was used to extract quantitative inhibition zones from the image, Supplementary Figure 3. The image was flattened using a gradient mask in Adobe Photoshop to remove illumination differences between the centre and edges of the plate. Then the image was thresholded to convert from grey scale to black and white. A mutational close was performed to remove graininess from the image. A Gaussian blur then applied to remove blank areas at the centre of inhibition zones left by the pipette tip. Finally, a threshold was applied and the pixels within each inhibition zone counted using *Analyze Particles* function in Fiji [34].

### Characterisation of Genetic Circuits

All plasmids were transformed into *E. coli* MG1655 by electroporation for characterisation. Overnight cultures of the relevant strains were grown in supplemented M9 media with the appropriate antibiotics in a shaking incubator at 37°C and 200 rpm. After 16 hours of growth, the cultures were diluted 1:1000 into fresh supplemented M9 media with antibiotics and grown for 6 hours in a shaking incubator at 37° C and 200 rpm. After 6 hours the cultures were diluted 1:100 into fresh supplemented M9 media with antibiotics and induced with the relevant concentration of inducer. 200 μL of each induced culture was then pipetted into a clean 96 well microtitre plate and sealed with a Breathe-Easy sealing membrane. It was incubated at 37° C and had constant double orbital shaking for 16 hours.

The microtitre plate was then removed and 1 μL of culture from each well was used to inoculate 200 μL of PBS in a clean round-bottom 96 well microtitre plate. Flow cytometry was performed on an Attune NxT Acoustic Focusing Cytometer with Attune NxT Autosampler (Thermo Fisher Scientific, UK). The Attune NxT Autosampler was set sample 20 μL from each well with 2 mixing cycles and 4 rinses between each sample. Forward and side scatter height and area measurements were always recorded. Height and area measurements in the appropriate fluorescent channels were also recorded. The fluorescence channels are detailed in Table 3. The flow cytometry data was processed using flopR [33] to remove debris and doublets.

**Table 3:**
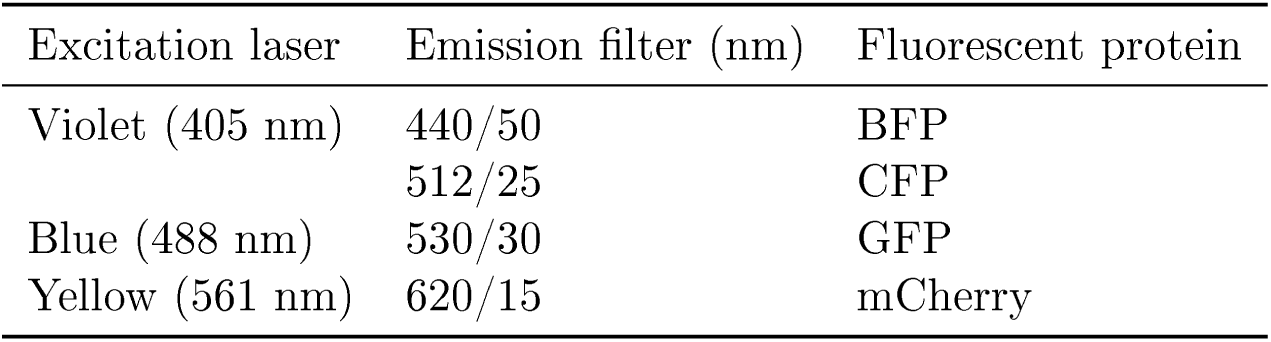
Fluorescence channels for the Attune NxT flow cytometer.

## Acknowledgements

We thank Y. Kaznessis and K. Geldart for plasmid pMPES:V, V. de Lorenzo and E. Martinez-Garcia for all SEVA plasmids and J. Ward for the E. coli MG1655 strain. We also thank N. Collant for guidance during some experiments.

## Author contributions

Conceptualization, AJHF and CPB; Formal Analysis, AJHF and BDK; Investigation, AJHF and BDK; Methodology, AJHF, BDK, MS and CPB; Project administration, CPB; Software, AJHF and BDK; Supervision, MS and CPB; Writing – original draft, AJHF and BDK; Writing - review & editing, AJHF, BDK, MS and CPB

## Data and materials availability

Plasmid maps and experimental data are available online.

