## Supplementary Information for "Single strain control of microbial consortia"

#### 1 Simple Model of Competition with Bacteriocins in Chemostat

We have developed a simple model of competition within a chemostat environment and added the ability of the engineered strain to produce a bacteriocin that can kill a competitor

$$
\begin{aligned}
\frac{dE}{dt} &= (\mu_E - D)E \\
\frac{dC}{dt} &= (\mu_C - D - k_\omega\, B)C \\
\frac{dS}{dt} &= D(S_0 - S) - \frac{\mu_E E}{\gamma_E} - \frac{\mu_C C}{\gamma_C} \\
\frac{dB}{dt} &= k_B E - DB
\end{aligned}
$$

where $E$ is the engineered population, $C$ is the competitor population, $S$ is the common growth substrate and $B$ is the bacteriocin, as shown in Figure 2, and the parameters are as defined in Table 1. The growth rates are given by Monod growth functions

$$\mu_E = \frac{\mu_{E_{\max}} S}{K_E + S}$$

$$\mu_C = \frac{\mu_{C_{\max}} S}{K_C + S}$$

The model is extended to include quorum molecule control of bacteriocin expression by the addition of a quorum molecule species, $Q$

$$\frac{dQ}{dt} = k_Q E - DQ$$

and making the production rate of the bacteriocin $k_B$ dependent on $Q$

$$k_B = (k_{B_{\max}} - k_{B_{\min}}) \frac{Q^n}{K_Q^n + Q^n} + k_{B_{\min}}$$

### 2 Steady State Analysis of the Simple Model

We make the simplifying assumption that $\gamma_E = \gamma_C = \gamma$ and $K_E = K_C = K_S$ i.e. both strains will utilise substrate in the same way, the only difference being the maximum rate at which each strain can grow. The model without the quorum molecule has four solutions at steady state

$$E = 0$$

$$C = 0$$

$$S = S_0$$

$$B = 0$$

Table 1: Model parameters

| Parameter | Description | Unit |
|---|---|---|
| $D$ | Dilution rate | $h^{-1}$ |
| $\mu_{E_{\max}}$ | Maximum growth rate of engineered strain | $h^{-1}$ |
| $\mu_{C_{\max}}$ | Maximum growth rate of competitor strain | $h^{-1}$ |
| $S_0$ | Reservoir substrate concentration | g $L^{-1}$ |
| $\gamma_E$ | Yield coefficient of engineered strain | cell $g^{-1}$ |
| $\gamma_C$ | Yield coefficient of competitor strain | cell $g^{-1}$ |
| $K_E$ | Concentration of substrate at which engineered strain growth is half-maximal | g $L^{-1}$ |
| $K_C$ | Concentration of substrate at which competitor strain growth is half-maximal | g $L^{-1}$ |
| $k_\omega$ | Death rate of competitor cells from bacteriocin | $M^{-1}$ $h^{-1}$ |
| $k_B$ | Expression rate of bacteriocin from engineered strain | mol $cell^{-1}$ $h^{-1}$ |
| $k_Q$ | Expression rate of quorum molecule from engineered strain | mol $cell^{-1}$ $h^{-1}$ |
| $k_{B_{max}}$ | Maximal expression rate of bacteriocin from engineered strain | mol $cell^{-1}$ $h^{-1}$ |
| $k_{B_{min}}$ | Minimal expression rate of bacteriocin from engineered strain | mol $cell^{-1}$ $h^{-1}$ |
| $K_Q$ | Concentration of $Q$ at which bacteriocin expression is half-maximal | M |
| $n$ | Cooperativity of bacteriocin repression by $Q$ | |

$$
\begin{aligned}
E &= \gamma(\frac{DK_S}{D-\mu_{E_{\max}}}+S_0) \\
C &= 0 \\
S &= \frac{DK_S}{\mu_{E_{\max}}-D} \\
B &= \gamma K_B(\frac{DK_S}{D-\mu_{E_{\max}}}+\frac{S_0}{D})
\end{aligned}
$$

$$
\begin{aligned}
E &= 0 \\
C &= \gamma(\frac{DK_S}{D-\mu_{C_{\max}}}+S_0) \\
S &= \frac{DK_S}{\mu_{C_{\max}}-D} \\
B &= 0
\end{aligned}
$$

$$
\begin{aligned}
E &= \frac{D^2(\mu_{C_{\max}}-\mu_{E_{\max}})}{k_B k_\omega \mu_{E_{\max}}} \\
C &= \frac{\frac{D(\gamma k_B k_\omega K_S \mu_{E_{\max}}+D^2(\mu_{E_{\max}}-\mu_{C_{\max}})+D\mu_{E_{\max}}(\mu_{C_{\max}}-\mu_{E_{\max}}))}{k_B k_\omega (D-\mu_{E_{\max}})}+\gamma\mu_{E_{\max}}S_0}{\mu_{C_{\max}}} \\
S &= \frac{DK_S}{\mu_{E_{\max}}-D} \\
B &= \frac{D(\mu_{C_{\max}}-\mu_{E_{\max}})}{k_\omega \mu_{E_{\max}}}
\end{aligned}
$$

The introduction of the quorum molecule makes the system of equations too complex to solve analytically. Instead, for each set of parameters, we simulate the system for 5000 hours and take the species at the final timepoint as the steady state values.

### 3 Model space exploration equations

$$
\begin{aligned}
\frac{dN_E}{dt} &= N_E(\mu_E - \eta_E - \omega_E - D) \\
\frac{dN_C}{dt} &= N_C(\mu_C - \omega_C - D) \\
\frac{dS}{dt} &= D(S_{0_j} - S_j) - \frac{N_E\mu_E}{\gamma} - \frac{N_C\mu_C}{\gamma} \\
\frac{dQ}{dt} &= k_Q N_E - DQ \\
\frac{dB}{dt} &= k_B(Q)N_E - DB \\
\frac{dI}{dt} &= k_I(Q)N_E - I\frac{\mu_E}{2} \\
\frac{dT}{dt} &= k_T(Q)N_E - T\frac{\mu_E}{2} - \alpha(T,V) \\
\frac{dV}{dt} &= k_V(Q)N_E - V\frac{\mu_E}{2} - \alpha(T,V)
\end{aligned}
$$

$$\omega_E = \begin{cases} 0 & \text{if } I \text{ constitutive,} \\ \omega_{\max} \frac{B^{n_\omega}}{K_\omega^{n_\omega} + B^{n_\omega}} \cdot \frac{K_I^{n_I}}{K_I^{n_I} + I^{n_I}} & \text{else} \end{cases}$$

$$\omega_C = \omega_{\max} \frac{B^{n_\omega}}{K_\omega^{n_\omega} + B^{n_\omega}}$$

$$\eta = \eta_{\max} \frac{T^{n_T}}{K_\eta^{n_\eta} + T^{n_\eta}}$$

$$\mu_E = \mu_{E_{\max}} \frac{S}{K_E + S}$$

$$\mu_C = \mu_{C_{\max}} \frac{S}{K_C + S}$$

$$k_B(Q) = \begin{cases} k_{B_{\max}} \frac{Q^{n_B}}{K_B^{n_B} + Q^{n_B}} & \text{if } B \text{ induced,} \\ k_{B_{\max}} \frac{K_B^{n_B}}{K_B^{n_B} + Q^{n_B}} & \text{if } B \text{ repressed,} \\ k_{B_{\max}} & \text{if } B \text{ constitutive,} \\ 0, & \text{if } B \text{ not expressed,} \end{cases} \qquad k_T(Q) = \begin{cases} k_{T_{\max}} \frac{Q^{n_T}}{K_T^{n_T} + Q^{n_T}} & \text{if } T \text{ induced,} \\ k_{T_{\max}} \frac{K_T^{n_T}}{K_T^{n_T} + Q^{n_T}} & \text{if } T \text{ repressed,} \\ k_{T_{\max}} & \text{if } T \text{ constitutive,} \\ 0 & \text{if } T \text{ not expressed,} \end{cases}$$

$$k_V(Q) = \begin{cases} k_{V_{\max}} \frac{Q^{n_V}}{K_V^{n_V} + Q^{n_V}} & \text{if } V \text{ induced,} \\ k_{V_{\max}} \frac{K_V^{n_V}}{K_V^{n_V} + Q^{n_V}} & \text{if } V \text{ repressed,} \\ 0 & \text{if } V \text{ not expressed,} \end{cases} \qquad k_I(Q) = \begin{cases} k_{I_{\max}} \frac{Q^{n_V}}{K_I^{n_I} + Q^{n_I}} & \text{if } I \text{ induced,} \\ k_{I_{\max}} \frac{K_I^{n_T}}{K_I^{n_I} + Q^{n_I}} & \text{if } I \text{ repressed,} \\ k_{I_{\max}} & \text{if } I \text{ constitutive,} \\ 0 & \text{if } I \text{ not expressed,} \end{cases}$$

#### Growth Equations

The change in the population of the engineered strain ($N_E$) is determined by the concentration of nutrient ($S$), bacteriocin ($B$), immunity protein ($I$), toxin proteins ($T$). The change in population of the competitor strain ($N_C$) is dependent upon the concentration of substrate and bacteriocin.

$$\mu_E = \mu_{E_{\max}} \frac{S}{K_E + S}$$
$$\mu_C = \mu_{C_{\max}} \frac{S}{K_C + S}$$

Detrimental effect of $B$ and the protection provided by $I$ are described using hill equations, whereby the negative effect of $B$ is repressed by $I$. If the engineered strain is constitutively expressing $I$, immunity is constant, therefore, $\omega_E$ goes to 0.

$$\omega_E = \begin{cases} 0 & \text{if I constitutive,} \\ \omega_{\max} \frac{B^{n_\omega}}{K_\omega^{n_\omega} + B^{n_\omega}} \cdot \frac{K_I^{n_I}}{K_I^{n_I} + I^{n_I}} & \text{else} \end{cases}$$

$$\omega_C = \omega_{\max} \frac{B^{n_\omega}}{K_\omega^{n_\omega} + B^{n_\omega}}$$

The detrimental effect of $T$ is also described as a hill function.

$$\eta = \eta_{\max} \frac{T^{n_T}}{K_\eta^{n_\eta} + T^{n_\eta}}$$

Bringing these functions together we can assemble differential equations describing the change in the population of a strain. The positive term being the growth rate, and negative terms being dilution rate, killing by bacteriocin and killing by toxin.

$$\frac{dN_E}{dt} = N_E(\mu_E - \eta - \omega_E - D)$$
$$\frac{dN_C}{dt} = N_C(\mu_C - \omega_C - D)$$

#### Nutrient species

The change in the concentration of $S$ is dependent on consumption by $N_E$ and $N_C$, and dilution with fresh media. $N_E$ and $N_C$ consume $S$ with a yield constant, $\gamma$. The dilution term describes the continuous replenishment of nutrient with the stock nutrient ($S_0$).

$$S = D(S_0 - S) - N_E\frac{\mu_E}{\gamma} - N_C\frac{\mu_C}{\gamma}$$

#### Expressed species

Quorum species, $Q$, is expressed at a rate that is linearly proportional to the engineered strain.

$$\frac{dQ}{dt} = k_Q N_E - DQ$$

The change in concentration of $B$ is effected by expression from strains in the system and dilution by the chemostat environment. The form taken by $k_B(Q)$ is dependent on the method of regulation, which varies between models.

$$\frac{dB}{dt} = k_B(Q)N_E - DB, \qquad k_B(Q) = \begin{cases} k_{B_{\max}}\frac{Q^{n_B}}{K_B^{n_B}+Q^{n_B}} & \text{if } B \text{ induced,} \\ k_{B_{\max}}\frac{K_B^{n_B}}{K_B^{n_B}+Q^{n_B}} & \text{if } B \text{ repressed,} \\ k_{B_{\max}} & \text{if } B \text{ constitutive,} \\ 0, & \text{if } B \text{ not expressed,} \end{cases}$$

Toxin ($T$), antitoxin ($V$) and immunity ($I$) species are all intracellular. The environmental dilution term is replaced with an intracellular dilution term derived from the rate of cell division of the engineered strain, $\frac{\mu_E}{2}$. Toxin and antitoxin species sequester one another, this is modelled as a linear relationship based on the concentration of the two species. Each species has different modes of expression which varies between the models. Constitutive expression of protective species $V$ gives complete protection from $T$, we therefore discard these models (see Model space definition). Constitutive expression of $I$ is reflected by changing $\omega_E$ to $0$.

$$\frac{dI}{dt} = k_I(Q)N_E - I\frac{\mu_E}{2}, \qquad k_I(Q) = \begin{cases} k_{I_{\max}}\frac{Q^{n_V}}{K_I^{n_I}+Q^{n_I}} & \text{if } I \text{ induced,} \\ k_{I_{\max}}\frac{K_I^{n_T}}{K_I^{n_I}+Q^{n_I}} & \text{if } I \text{ repressed,} \\ 0 & \text{if } I \text{ not expressed,} \end{cases}$$

$$\frac{dT}{dt} = k_T(Q)N_E - T\frac{\mu_E}{2} - \alpha(T,V), \quad k_T(Q) = \begin{cases} k_{T_{\max}}\frac{Q^{n_T}}{K_T^{n_T}+Q^{n_T}} & \text{if } T \text{ induced,} \\ k_{T_{\max}}\frac{K_T^{n_T}}{K_T^{n_T}+Q^{n_T}} & \text{if } T \text{ repressed,} \\ k_{T_{\max}} & \text{if T constitutive,} \\ 0 & \text{if } T \text{ not expressed,} \end{cases}$$

$$\frac{dV}{dt} = k_V(Q)N_E - V\frac{\mu_E}{2} - \alpha(T,V), \quad k_V(Q) = \begin{cases} k_{V_{\max}}\frac{Q^{n_V}}{K_V^{n_V}+Q^{n_V}} & \text{if } V \text{ induced,} \\ k_{V_{\max}}\frac{K_V^{n_V}}{K_V^{n_V}+Q^{n_V}} & \text{if } V \text{ repressed,} \\ 0 & \text{if } V \text{ not expressed,} \end{cases}$$

$$\alpha(T,V) = k_{\text{ann}} \cdot T \cdot V$$

#### Example system - Figure 4Div

The engineered strain constitutively expresses $Q$. $B$ is repressed by $Q$.

$$\begin{aligned} \frac{dN_E}{dt} &= N_E(\mu_E - D) \\ \frac{dN_C}{dt} &= N_C(\mu_C - \omega_C - D) \\ \frac{dS}{dt} &= D(S_{0_j} - S_j) - \frac{N_E\mu_E}{\gamma} - \frac{N_C\mu_C}{\gamma} \\ \frac{dQ_y}{dt} &= k_Q N_E - DQ \\ \frac{dB}{dt} &= \frac{K_B^{n_B}}{K_B^{n_B} + Q^{n_B}} N_E - DB \end{aligned}$$

#### Model space definition

Species $B$, $I$ and $T$ can be expressed constitutively, induced, repressed or not expressed. $Q$ can be either expressed constitutively or not expressed. $V$ can be induced, repressed or not expressed. We represent expression options using integers. $0$ constitutive, $1$ induced, $2$ repressed, $3$ not expressed. We can define the options for each expressed species as such.

$$B = 0 \text{ or } 1 \text{ or } 2 \text{ or } 3$$

$$I = 0 \text{ or } 1 \text{ or } 2 \text{ or } 3$$

$$T = 0 \text{ or } 1 \text{ or } 2 \text{ or } 3$$

$$V = 1 \text{ or } 2 \text{ or } 3$$

$$B = 0 \text{ or } 3$$

We restrict the engineered strain to express up to one of each Let $P_c$ be the number of unique part combinations, choosing one expression option for the engineered strain in each model

$$P_C = \binom{B}{1} \times \binom{I}{1} \times \binom{T}{1} \times \binom{V}{1} \times \binom{Q}{1}$$

We use a series of rules to remove illegal and redundant models from the model space.

Combination is illegal if any of $B$, $I$, $T$ or $V$ are regulated by $Q$, while $Q$ is not expressed

$$C_1 : (B \text{ or } I \text{ or } T \text{ or} V_i \in \{1, 2\} \text{ and } Q_i = 0)$$

Combination is illegal if $I$ is expressed, and $B$ is not expressed

$$C_2 : (I_i \in \{0, 1, 2\} \to B_i = 3)$$

Combination is illegal if $V$ is expressed and $T$ is not expressed

$$C_3 : (V_i \in \{1, 2\} \rightarrow T_i = 3)$$

Combination is illegal when $Q$ is expressed and none of $B$, $I$, $T$ or $V$ are regulated by $Q$

$$C_5 : (Q_i = 0 \rightarrow B_i \ \& \ I_i \ \& \ T_i \ \& \ V_i \notin \{1, 2\})$$

Legal part combinations are taken by substracting illegal combinations from the total unique part combinations. $P_L$ is a set containing all the different ways in which the engineered strain can be engineered.

$$P_L = P_C - (C_1 \cup C_2 \cup C_3 \cup C_4 \cup C_5)$$

Table 2: Model priors

| Parameter | Description | Lower bound | Upper bound | Prior distribution | Unit |
|---|---|---|---|---|---|
| $D$ | Dilution rate | $0.01$ | $0.5$ | Uniform | $h^{-1}$ |
| $\mu_{E_{\max}}$ | Maximum growth rate of engineered strain | $0.4$ | $3$ | Uniform | $h^{-1}$ |
| $\mu_{C_{\max}}$ | Maximum growth rate of competitor strain | $0.4$ | $3$ | Uniform | $h^{-1}$ |
| $\gamma$ | Yield coefficient | $1e^{12}$ | $1e^{12}$ | Constant | cell $g^{-1}$ |
| $S_0$ | Concentration of $S$ in input media | $4$ | $4$ | Constant | g $L^{-1}$ |
| $\omega_{\max}$ | Maximal death rate due to bacteriocin | $0.5$ | $2$ | Uniform | $h^{-1}$ |
| $\eta_{\max}$ | Maximal death rate due to bacteriocin | $0.5$ | $2$ | Uniform | $h^{-1}$ |
| $K_\omega$ | Concentration of $B$ at which killing rate is half-maximal | $1e^{-12}$ | $1e^{-7}$ | Uniform | $M^{-1}$ |
| $n_\omega$ | Cooperativity coefficient for detrimental effect of bacteriocin | $1$ | $2$ | Uniform | |
| $K_E$ | Concentration of $S$ at which growth of engineered strain is half-maximal | $2$ | $2$ | Constant | $gL^{-1}$ |
| $K_C$ | Concentration of $S$ at which growth of engineered strain is half-maximal | $2$ | $2$ | Constant | $gL^{-1}$ |
| $K_B$ | Concentration of $Q$ at which expression of bacteriocin is half-maximal | $1e^{-9}$ | $1e^{-7}$ | Log uniform | M |
| $K_I$ | Concentration of $Q$ at which expression of immunity is half-maximal | $1e^{-9}$ | $1e^{-7}$ | Log uniform | M |
| $K_T$ | Concentration of $Q$ at which expression of toxin is half-maximal | $1e^{-9}$ | $1e^{-7}$ | Log uniform | M |
| $K_V$ | Concentration of $Q$ at which expression of anti-toxin is half-maximal | $1e^{-9}$ | $1e^{-7}$ | Log uniform | M |
| $K_\eta$ | Concentration of $T$ at which repression of growth is half-maximal | $1e^{-21}$ | $1e^{-16}$ | Log uniform | M |
| $n_\eta$ | Cooperativity coefficient for detrimental effect of toxin | $1$ | $2$ | Uniform | $M^{-1}h^{-1}$ |
| $k_{B_{\max}}$ | Maximum expression rate of bacteriocin | $1e^{-21}$ | $1e^{-19}$ | Log uniform | $M^{-1}h^{-1}$ |
| $k_{I_{\max}}$ | Maximum expression rate of immunity | $1e^{-21}$ | $1e^{-19}$ | Log uniform | $M^{-1}h^{-1}$ |
| $k_{T_{\max}}$ | Maximum expression rate of toxin | $1e^{-21}$ | $1e^{-19}$ | Log uniform | $M^{-1}h^{-1}$ |
| $k_{V_{\max}}$ | Maximum expression rate of anti-toxin | $1e^{-21}$ | $1e^{-19}$ | Log uniform | $M^{-1}h^{-1}$ |
| $n_B$ | Cooperativity coefficient for expression of bacteriocin | $2$ | $2$ | Constant | |
| $n_I$ | Cooperativity coefficient for expression of immunity | $2$ | $2$ | Constant | |
| $n_T$ | Cooperativity coefficient for expression of toxin | $2$ | $2$ | Constant | |
| $n_V$ | Cooperativity coefficient for expression of anti-toxin | $2$ | $2$ | Constant | |
| $k_Q$ | Production rate of $Q$ | $1e^{-20}$ | $1e^{-19}$ | Log uniform | $M^{-1}h^{-1}$ |
| $k_{ann}$ | Rate of toxin anti-toxin annihilation | $30$ | $30$ | Constant | $M^{-1}h^{-1}$ |

### 4 Model selection with ABC SMC

#### Bayesian inference

Let $\theta \in \Theta$ be a parameter vector with a prior $\pi(\theta)$. Given an objective of $x_0$, where $x_0$ is exists in the solution space, $x_0 \in \mathcal{D}$. We define the likelihood function for the objective behaviour as $f(x_0|\theta)$.

$$\pi(\theta|x_0) = \frac{f(x_0|\theta)\pi(\theta)}{\pi(x_0)}$$

We can rewrite $\pi(x_0)$ where $a$ and $b$ represent the lower and upper bounds of the parameter value:

$$\pi(x) = \int_a^b f(x_0, \theta)d\theta = \int_a^b f(x_0|\theta)\pi(\theta)d\theta$$

The posterior distribution informs us of the parameter distribution that gives rise to the objective.

$$\pi(\theta|x_0) = \frac{f(x_0|\theta)\pi(\theta)}{\int_a^b f(x_0|\theta)\pi(\theta)d\theta}$$

Let $m$ be a model from vector of competing models, $M$,such that $m \in M = \{m_1, m_2 ... m_q\}$. Each model has its own parameter space, allowing us to define a joint space, $(m, \theta) \in M \times \Theta_M$.

We can write Bayes' theorem in terms from the context of a model space.

$$\pi(m|x_0) = \frac{f(x_0|m)\pi(m)}{\int_M f(x_0|m')\pi(m')dm'}$$

Since the $M$ is discrete, we can rewrite this

$$\pi(m|x_0) = \frac{f(x_0|m)\pi(m)}{\sum_M f(x_0|m')\pi(m')}$$

The marginal likelihood of the model, $f(x_0|m)$, considers the joint parameter and model space

$$f(x_0|m) = \int_{\Theta M} \pi(\theta|m) f(x_0|\theta, m)d\theta$$

#### Approximate Bayesian computation

In our case an analytical solution of the likelihood function, $f(x_0|\theta)$, cannot be derived because our objective behaviour has an infinite number of possible solutions. We bypass this and approximate the posterior by generating data from a model. We can sample a parameter vector from the prior, $\theta^* \sim \pi(\theta)$, which is simulated to yield a data vector, $x^*$. This can be written as a conditional, $x^* \sim f(x|\theta^*)$, which also gives the joint density, $\pi(\theta, x)$.

In order to obtain the posterior distribution that satisfies our objective behaviour, $x_0$, we apply a conditional to define whether a generated data vector, $x^*$ belongs to the objective $x_0$.

Let us set an indicator function $\mathbb{I}$

$$\mathbb{I}(x) = \begin{cases} 1, & \text{if } x = x_0 \\ 0, & \text{otherwise} \end{cases}$$

The indicator function will be $1$ if the simulation data has the required characteristics of the objective, and $0$ if it is not. By applying this condition we have generated a rejection algorithm that allows us to obtain the posterior distribution.

$$\pi(\theta|x, x_0) = \frac{\pi(\theta)f(x|\theta)\mathbb{I}(x)}{\int_{\mathcal{A}_{x_0}\times\Theta} \pi(\theta)f(x|\theta)dxd\theta}$$

We replace $\mathbb{I}_{\mathcal{A}_{x_0}}$ with $\mathbb{I}_{\mathcal{A}_{x_0}}(x, \epsilon)$. $\rho$ is function that outputs a distance between $x_0$ and $x$, telling us how far away the simulated data is to the objective data. $\epsilon$ defines the threshold below which the distance is acceptably small.

$$\mathbb{I}(x, \epsilon) = \begin{cases} 1, & \text{if } \rho(x, x_0) = \epsilon \\ 0, & \text{otherwise} \end{cases}$$

Giving us $\pi_\epsilon$, the approximation of the posterior

$$\pi_\epsilon(\theta|x, x_0) = \frac{\pi(\theta)f(x|\theta)\mathbb{I}(x)}{\int_{\mathbb{I}=1\times\Theta} \pi(\theta)f(x|\theta)dxd\theta}$$

The smaller $\epsilon$ is and the larger the number of simulations conducted, the more accurate the representation of the true posterior will be. We can write this marginal posterior distribution as

$$\pi(\theta^*|\rho(x^*, x_0)) \leq \epsilon \approx \pi(\theta|x_0)$$

#### ABC rejection algorithm

The most basic ABC algorithm is the ABC rejection algorithm. Let $\epsilon$ be the distance threshold the defining the necessary level of agreement between the objective, $x_0$, and a given simulation, $x^*$

**Algorithm 1:** ABC rejection algorithm

Set particle indicator $i = 0$
**while** $i < N$ **do**
    Sample model $m$, from model space prior, $M$
    Sample parameter $\theta^*$, from prior distribution $\pi(\theta)$
    Simulate model, $f(x|\theta^*)$, giving simulation data, $x^*$.
    Calculate distance between simulation data and objective, $d(x^*, x)$.
    **if** $d(x^*, x) \leq \epsilon$ **then**
        Set $\theta_t^i = \theta^*$
        i = i + 1
    **else**
        Reject $\theta^*$
**end while**
Posterior distribution generated from accepted particles, $\pi(\theta^*|d(x^*, x)) \leq \epsilon$:

#### Model selection with ABC SMC

In this paper we use a variant of ABC, ABC Sequential Monte Carlo (ABC SMC) (*1*). ABC SMC evolves the prior distribution through a series of intermediate distributions. The distance threshold ($\epsilon$) is decreased between distributions, moving the acceptance criteria closer to the objective. The gradual evolution reduces the tendency of becoming focused on local areas of minimal distance.

**Algorithm 2:** Model selection with ABC SMC

1 Set population indicator, $t = 0$
2 Set initial epsilon, $\epsilon_t = \inf$
3 Set final epsilon, $\epsilon_T = [x, y, z]$
4 Set particle indicator, $i = 0$
5 **if** $t = 0$ **then**
6 Sample $m^*$ from $\pi(m)$
7 Sample $\theta^{**}$ from $\pi(\theta(m^*))$
8 **else if** $t > 0$ **then**
9 Sample particle $\theta^*$ from previous population $\{\theta(m^*)^i_{t-1}\}$ with weights $w(m^*)_{t-1}$
10 Perturb $\theta^*$ to obtain $\theta^{**} \sim K_t(\theta|\theta^*)$
11 **if** $\pi(\theta^{**}) = 0$ **then**
12 **go to** 5
13
14 Simulate, $x^* \sim f(x|\theta^{**}, m^*)$
15 **if** $d(x^*, x_0) > \epsilon_t$ **then**
16 **go to** 5
17
18 Set $m^i_t = m^*$
19 Set $\theta^i_t = \theta^{**}$
20 Calculate particle weight, $w^i_t$
21 **if** $t = 0$ **then**
22 $w^i_t = 1$
23 **else**
24 $w^i_t = \frac{\pi(\theta^{**})}{\sum_{j=1}^{N} w^j_{t-1} K_t(\theta^i_{t-1}|\theta^{**})}$
25 **if** $i < N$ **then**
26 Set $i = i + 1$
27 **go to** 5
28
29 Normalise weights for every $m$.
30 **if** $\epsilon_t \neq \epsilon_T$ **then**
31 Update population number, $t = t + 1$
32 Update $\epsilon$ according to accepted particle distances, $\epsilon_t = f_\epsilon()$
33 **go to** 5

In these experiments we use a component-wise gaussian perturbation kernel. This produces a random walk from a particle of the previous population to a particle of the next population.

$$K_t(\theta|\theta^*) = \mathcal{N}(\theta^*, 2x) \tag{1}$$

Where, $x$, is the variance of the previous population.

$$x = \sigma(\theta_{t-1})^2$$

#### Defining stable steady state objective

We define the stable steady state objective with three summary statistics. Where $x$ is the time series data of a strain ($N_E$ or $N_C$). Each distance function has been chosen to filter out undesirable behaviours leaving the remaining desired stable steady state behaviour.

$d_1$ is the final gradient of $x$ is above a tolerance threshold

$$d_1(x) = |\Delta x(t-1)|$$

$d_2$ is the standard deviation of the signal, this filters out simulations which are oscillating.

$$d_2(x) = \sigma(x)$$

$d_3$ is the reciprocal the final value of the simulation, this allows us to define a minimum population threshold.

$$d_3(x) = \frac{1}{x(t-1)}$$

Using these distances we can define the final conditional $\epsilon_F$. The final threshold values we use are $\epsilon_F = \{500, 25000, 1e^{-10}\}$

$$d_1 < \epsilon_{F_1}$$

$$d_2 < \epsilon_{F_2}$$

$$d_3 < \epsilon_{F_3}$$

#### Auto $\epsilon$ generation

The next $\epsilon$ for each new population is generated based on the accepted particle distances from previous population until we reach the final $\epsilon$. We calculate distances for each species we are fitting to the objective ($N_E$ and $N_C$).

Let $X$ be time series data of a simulated particle, and $\hat{X}$ contain all accepted particles in a population, such that

$$X \in \hat{X}$$

From each simulation, we can take the time series data of the species to be fit, $X_{N_E}$ and $X_{N_C}$.

$$X_{N_E} \in X$$

$$X_{N_E} \in X$$

These time series data are used to calculate the distances of the particle from the objective. We couple the distances of the two species being fit by taking the maximum of the two for each distances.

$$V = \{X \in \hat{X} \mid \max(\{d_i(X_{N_E}), d_i(X_{N_C})\})\}$$

We set $\alpha$ as a proportion of the distances we should to generate the next $\epsilon$. Generally speaking, the samller $\alpha$ is the faster we will progress to $\epsilon_F$

$$n = [length(\hat{X})/\alpha]$$

Let $S$ be the $n$-th smallest distance, where $S \in V$. Allowing us to set the threshold for the next population, $\epsilon_{t+1}$.

$$\epsilon_{t+1} = S$$

### 5 Supplementary Figures

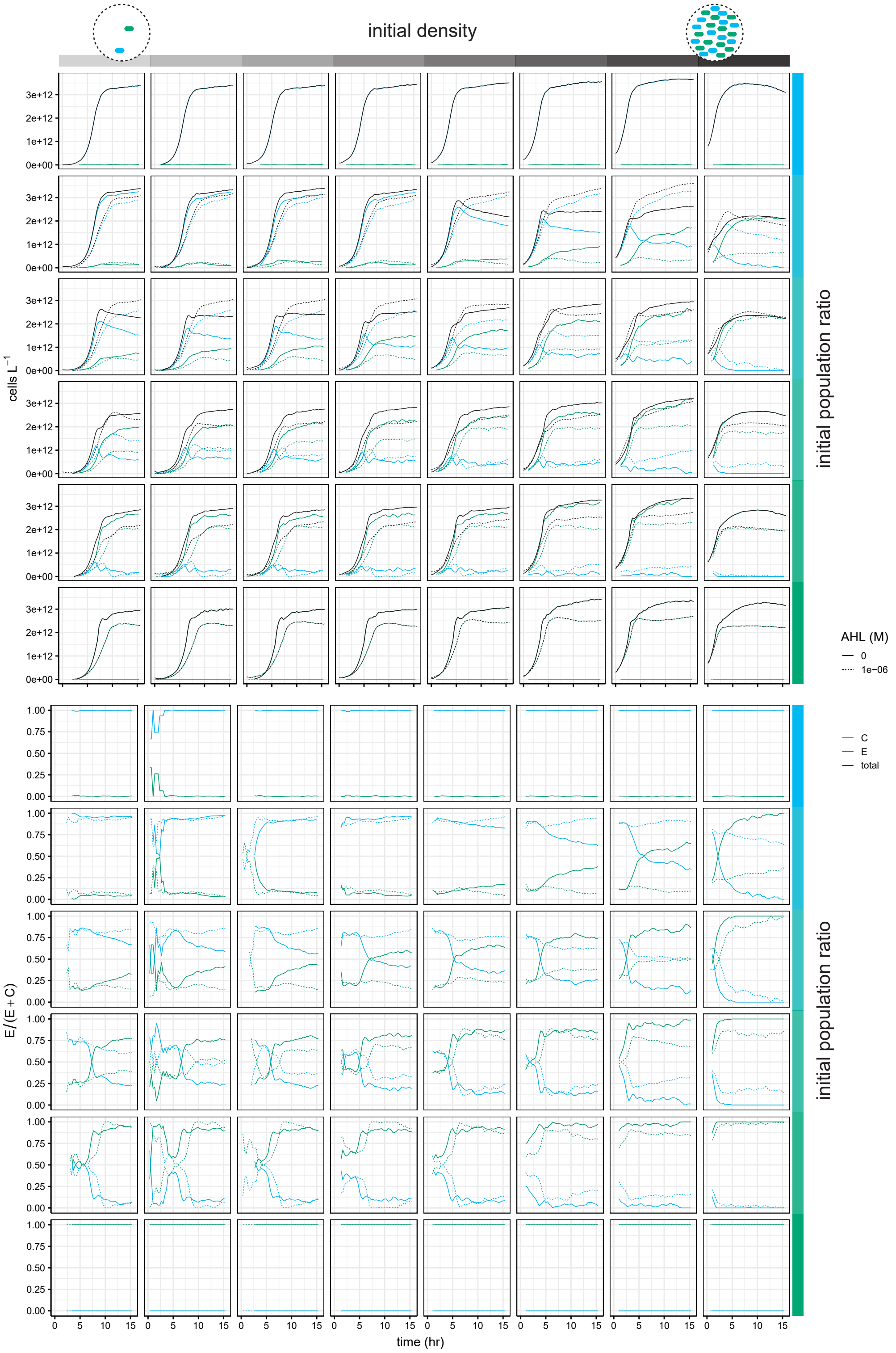


Figure 1: Competition at a variety of initial densities and population ratios. Solid lines show data used in Figure 1G.

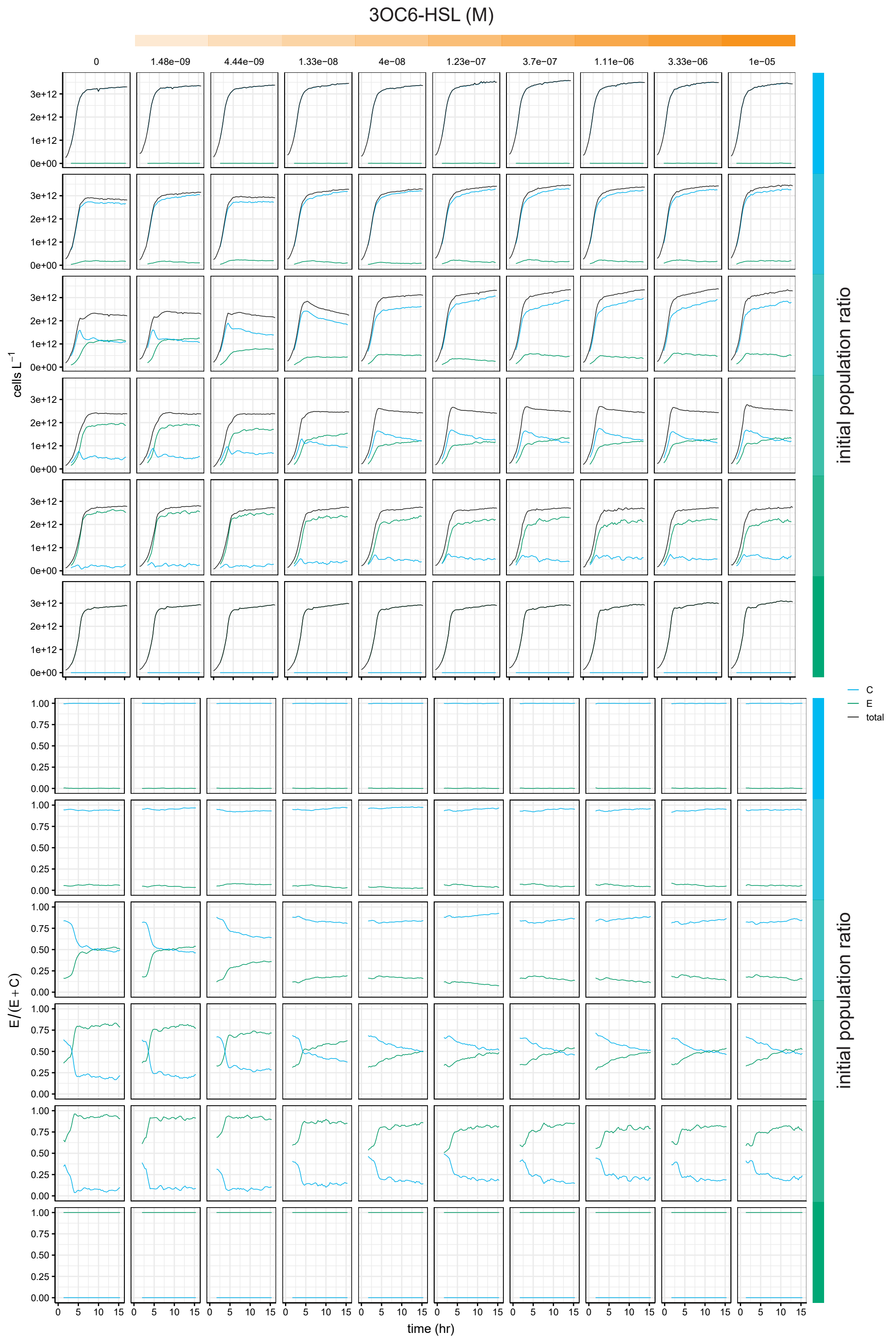


Figure 2: Competition at a variety of quorum molecule concentrations and initial population ratios used in Figure 2F.

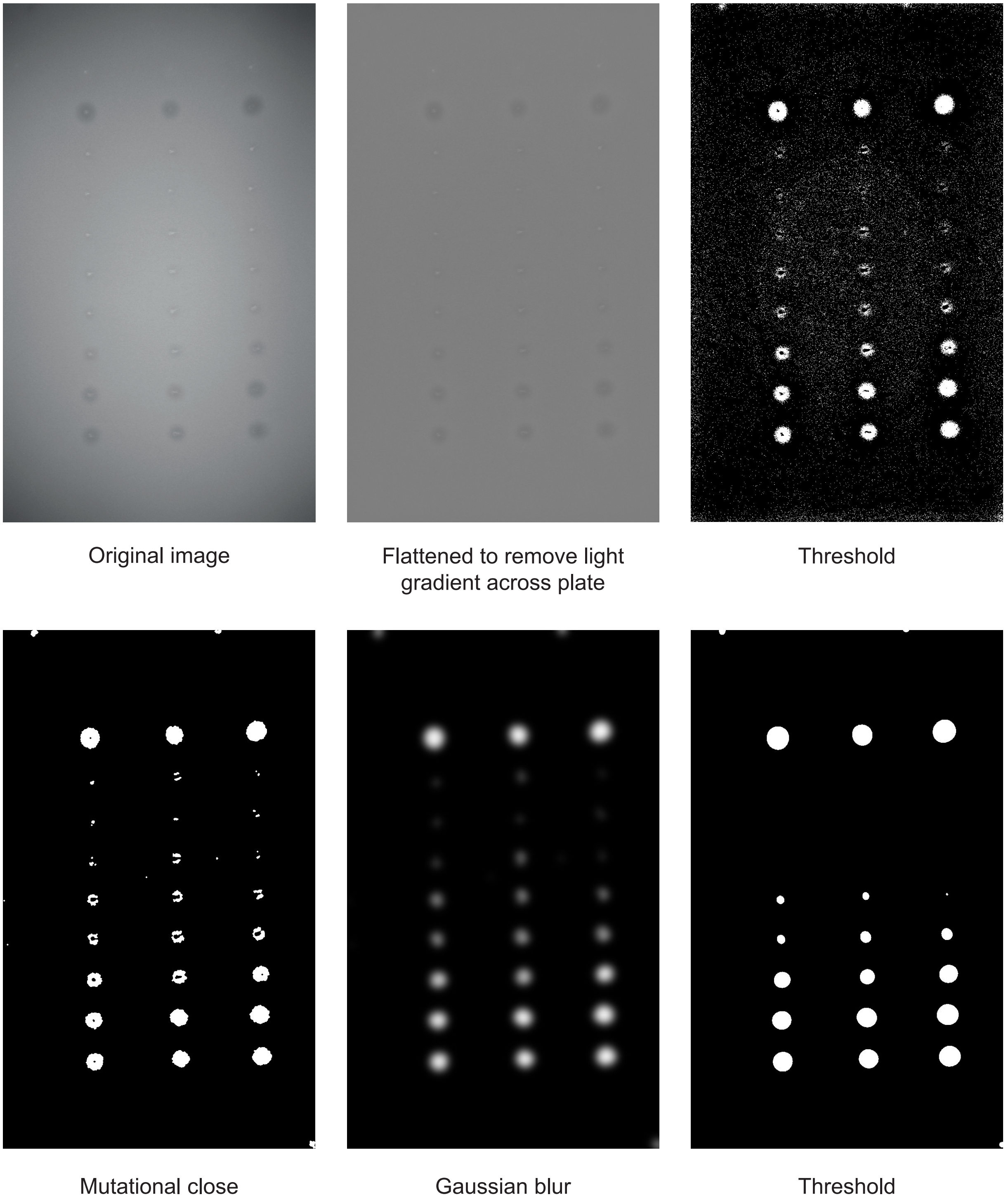


Figure 3: Image processing of agar plate spot inhibition assay.

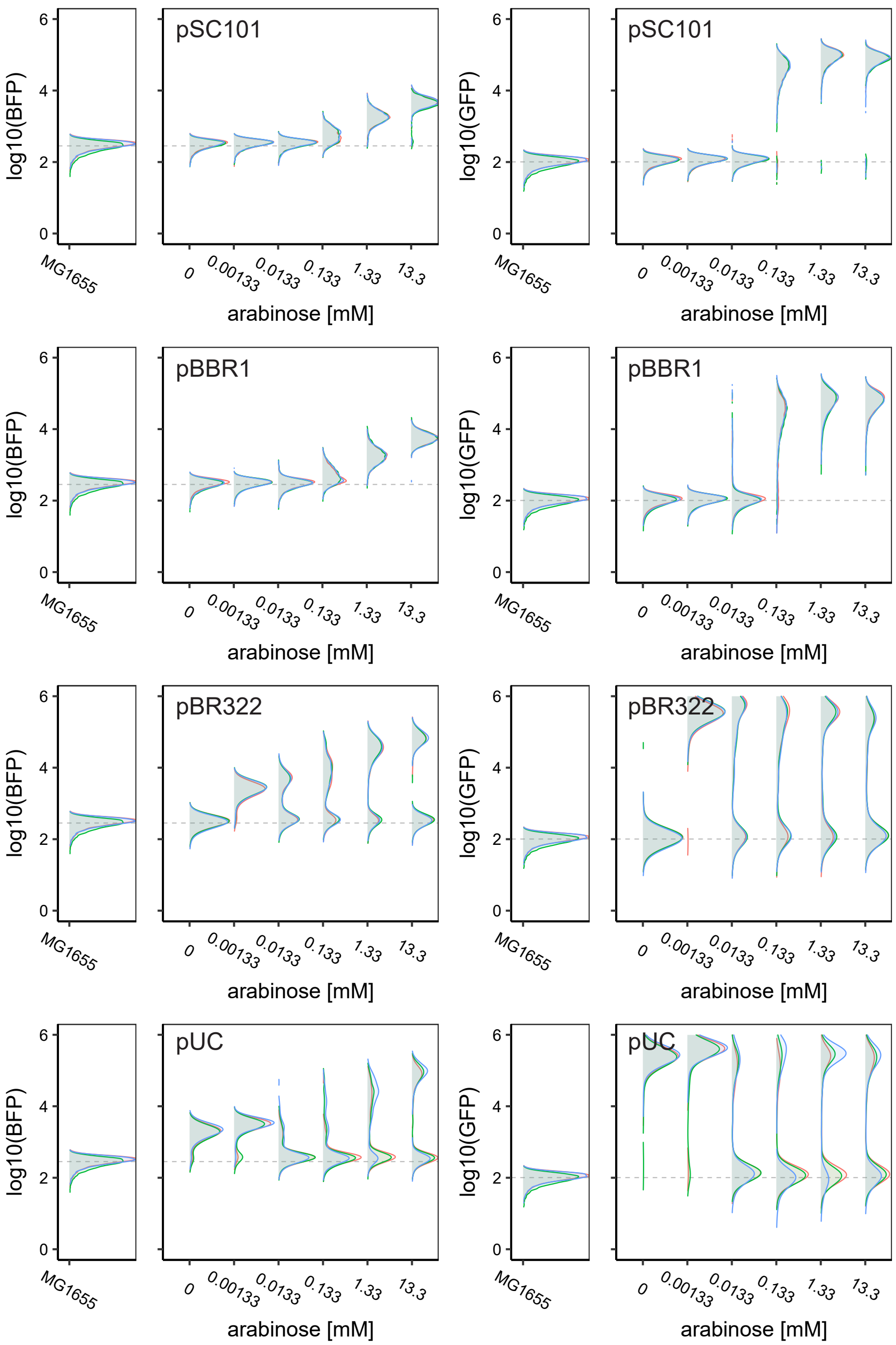


Figure 4: Changing plasmid copy number of the plasmid carrying the arabinose inducible LuxI (BFP) and 3OC6-HSL inducible TetR (GFP).

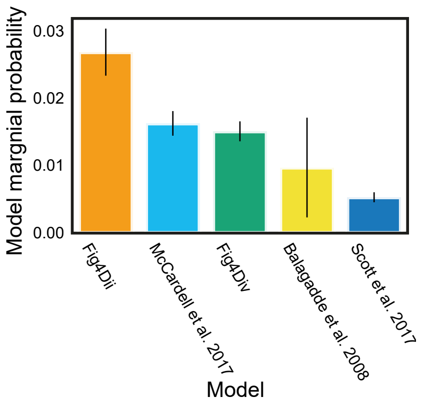


Figure 5: Comparison of model marginal probabilities, with a coexistence objective, for our system relative to previously described systems.
